# The role of animal faces in the animate-inanimate distinction in the ventral temporal cortex

**DOI:** 10.1101/2020.10.08.330639

**Authors:** D. Proklova, M.A. Goodale

**Author notes:** Correspondence. Address correspondence to Daria Proklova at, The Brain and Mind Institute, Western University, Western Interdisciplinary Research Building, London, ON N6A 3K7, Canada.

## Abstract

Animate and inanimate objects elicit distinct response patterns in the human ventral temporal cortex (VTC), but the exact features driving this distinction are still poorly understood. One prominent feature that distinguishes typical animals from inanimate objects and that could potentially explain the animate-inanimate distinction in the VTC is the presence of a face. In the current fMRI study, we investigated this possibility by creating a stimulus set that included animals with faces, faceless animals, and inanimate objects, carefully matched in order to minimize other visual differences. We used both searchlight-based and ROI-based representational similarity analysis (RSA) to test whether the presence of a face explains the animate-inanimate distinction in the VTC. The searchlight analysis revealed that when animals with faces were removed from the analysis, the animate-inanimate distinction almost disappeared. The ROI-based RSA revealed a similar pattern of results, but also showed that, even in the absence of faces, information about agency (a combination of animal’s ability to move and think) is present in parts of the VTC that are sensitive to animacy. Together, these analyses showed that animals with faces do elicit a stronger animate/inanimate response in the VTC, but that this effect is driven not by faces per se, or the visual features of faces, but by other factors that correlate with face presence, such as the capacity for self-movement and thought. In short, the VTC appears to treat the face as a proxy for agency, a ubiquitous feature of familiar animals.

**Significance Statement:** Many studies have shown that images of animals are processed differently from inanimate objects in the human brain, particularly in the ventral temporal cortex (VTC). However, what features drive this distinction remains unclear. One important feature that distinguishes many animals from inanimate objects is a face. Here, we used fMRI to test whether the animate/inanimate distinction is driven by the presence of faces. We found that the presence of faces did indeed boost activity related to animacy in the VTC. A more detailed analysis, however, revealed that it was the association between faces and other attributes such as the capacity for self-movement and thinking, not the faces *per se*, that was driving the activity we observed.

## Introduction

Multiple studies have shown that an animate/inanimate distinction is a major factor in the organization of object representations in the human ventral temporal cortex (VTC) (Kriegeskorte et al., 2008; Bracci and Op de Beeck, 2016; Proklova et al., 2016). Animacy, however, is associated with different attributes, from visual features to semantic concepts such as agency (Peelen and Downing, 2017), and it is still unclear what drives the animate/inanimate distinction in the VTC. Some studies have suggested that the it can be explained by visual features alone (Long et al., 2018; Coggan et al., 2016). Others have suggested that the animate/inanimate distinction is not based purely on visual features but is also driven by conceptual information associated with animacy (Thorat et al., 2019; Proklova et al., 2016; Sha et al., 2016).

Stimuli that do not conform to the strict animate/inanimate dichotomy have recently provided new insights into animacy organization in the VTC. A recent fMRI study showed that inanimate objects that share features with animals (e.g., cow-shaped mugs) are represented in a similar way to animate objects (Bracci et al., 2018). Another study that used “borderline” stimuli, such as robots, together with more typical animate or inanimate objects, found similar results using MEG (Contini et al., 2019). This suggests that what matters for the VTC is similarity to humans (either visual or semantic), rather than animacy *per se* (Contini et al., 2019; Sha et al., 2016; Gobbini et al., 2011; Thorat et al., 2019). Another possibility is that the VTC is tuned to the diagnostic features of animacy shared between animals and animal-like stimuli (e.g., faces or bodies; Bracci et al., 2018).

Although there is evidence that body shape cannot fully account for the representation of animacy in the VTC (Proklova et al., 2016, Bracci et al., 2016), few previous studies have controlled for the presence of a face. Faces are extremely biologically relevant stimuli. They play an important role in determining whether something is animate – and the animals with which people are most familiar tend to have faces. Animal-like objects such as robots and toys often share this critical feature with animals, which could explain why they are represented similarly to animals in the VTC.

A network of areas mediates face perception, most notably the fusiform face area (FFA) in the VTC (Grill-Spector et al., 2018). Interestingly, in a study by Proklova et al. (2016), the fusiform gyrus was one of the areas in which animacy information was present after controlling for most visual features. Since all the animate stimuli in that study had faces, it is possible that this effect was driven by the presence of faces rather than by animacy *per se*.

A number of neuroimaging studies have argued that faces are not necessary for eliciting typical animate/inanimate distinction in the VTC by using images of animals with their faces covered (Chao et al., 1999), geometric shapes moving in characteristically animate (or inanimate) ways (Martin and Weisberg, 2004), or synthetic stimuli (*texforms*) which preserve the mid-level features of the stimuli but not the fine detail such as faces (Long et al., 2018). To our knowledge, however, no previous studies have directly compared the patterns of brain activation generated by images of real animals with and without faces. In the present fMRI study, we addressed this gap by using a stimulus set that included images of real animals with and without a face, as well as inanimate objects, all of which had otherwise similar visual features. Thus, we made animacy orthogonal to the presence of a face, which allowed us to directly examine the role of faces in VTC representations. We found that when animals with faces were removed from the analysis, the animate/inanimate distinction largely disappeared. Nevertheless, additional analyses revealed that this effect was driven not by faces *per se*, but by other features that typically correlate with the presence of a face, such as mobility and thoughtfulness.

## Materials and methods

### Participants

Twenty-four volunteers (18 female, mean age 26.7 years, SD = 3.3) participated in the behavioral ratings experiment. These included 14 people who also took part in the fMRI experiment described below (rating the animals after the fMRI session) and ten who did not. Ten other volunteers, who were not part of the behavioral ratings or the fMRI experiments (4 female, mean age = 25 years, SD = 4.6) participated in the behavioral visual search experiment. Twenty volunteers (14 female, mean age 26.1 years, SD = 4) took part in the fMRI study. Two were excluded from further analysis because of excessive head movement. All participants gave informed consent and the protocol was approved by the University of Western Ontario Ethics Review Board.

### Stimuli

The stimulus set (Fig. 1) consisted of 18 unique stimuli divided into 3 groups: 6 animals with distinct faces (e.g., snake), 6 animals without a distinct face (e.g., starfish), and 6 inanimate objects. Additionally, we used three different exemplars of each stimulus, resulting in a total of 54 stimuli. Importantly, to minimize the effects of visual feature similarity, stimuli were organized into six triplets (animal with a face - faceless animal - object) based on the overall shape similarity, such that all three stimuli within each triplet shared a similar overall shape (e.g., snake – worm – rope).

**Figure 1.**
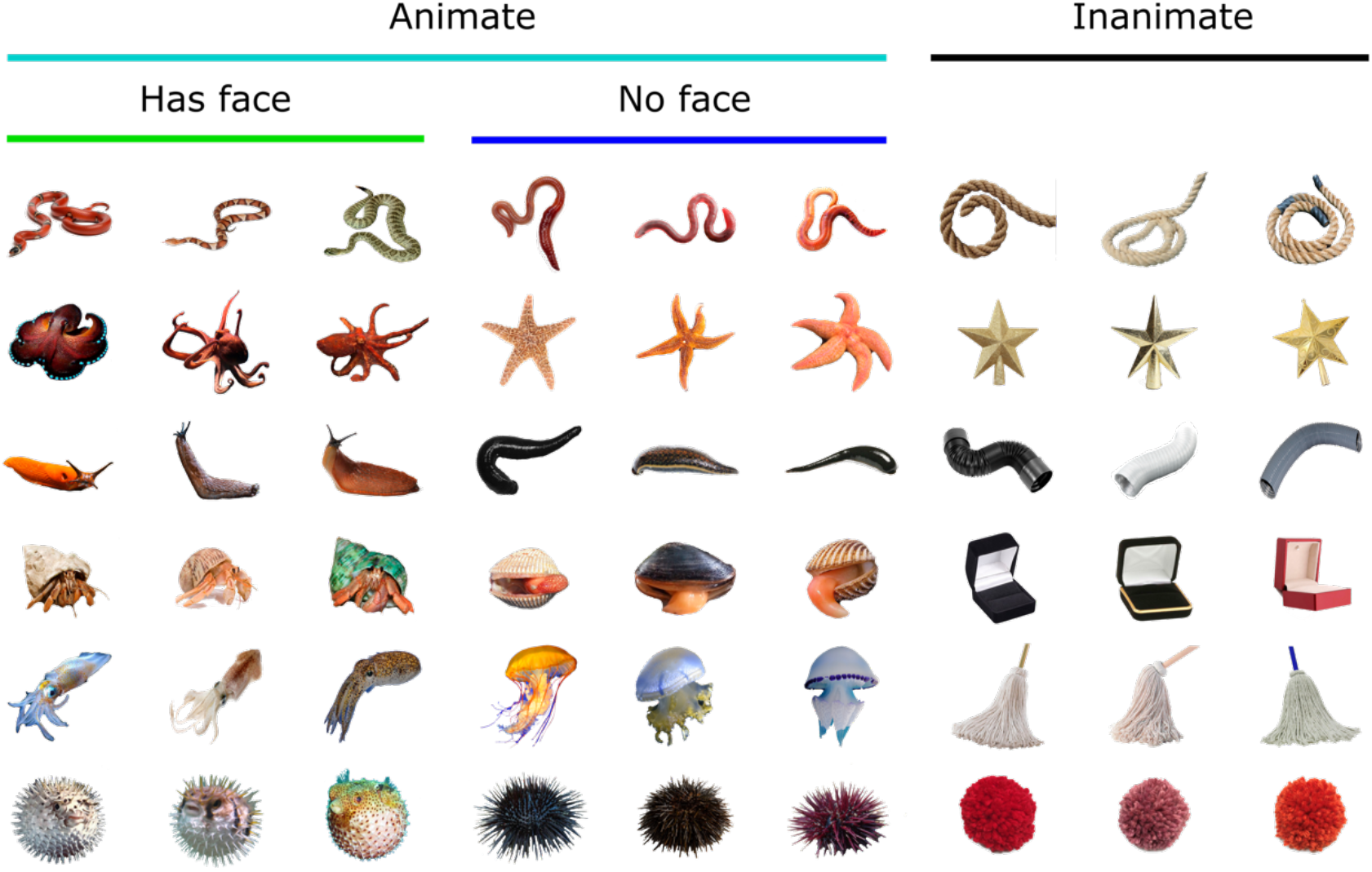
Stimulus set. The stimulus set included images of animals with faces (left 3 columns), without faces (middle 3 columns), and inanimate objects (right 3 columns). There were three exemplars of each unique animal/object. To minimize visual differences between the stimuli from different categories, the stimuli in each row shared a similar shape.

### Behavioral ratings experiment

Participants were asked to rate the 12 animal stimuli presented on a set of 6 characteristics: (1) Does this animal have a face? (2) How fast can this animal move? (3) Is this animal capable of thoughts? (4) Does this animal have a head? (5) Does this animal have eyes? (6) How familiar is this animal to you? There was a separate block for each of the 6 questions. The question appeared at the beginning of the block, followed by a presentation of all twelve animals used in the experiment. On each trial, participants were presented with all 3 versions of a given animal and the continuous ratings bar on the computer screen (Fig. 2C). Using a mouse, participants could click anywhere on a bar to provide their response, ranging from 0 (e.g. “this animal is not at all familiar”) to 100 (“I am very familiar with this animal”). In this and all the following experiments, including fMRI, stimuli were presented using Psychtoolbox for Matlab (Brainard, 1997).

**Figure 2.**
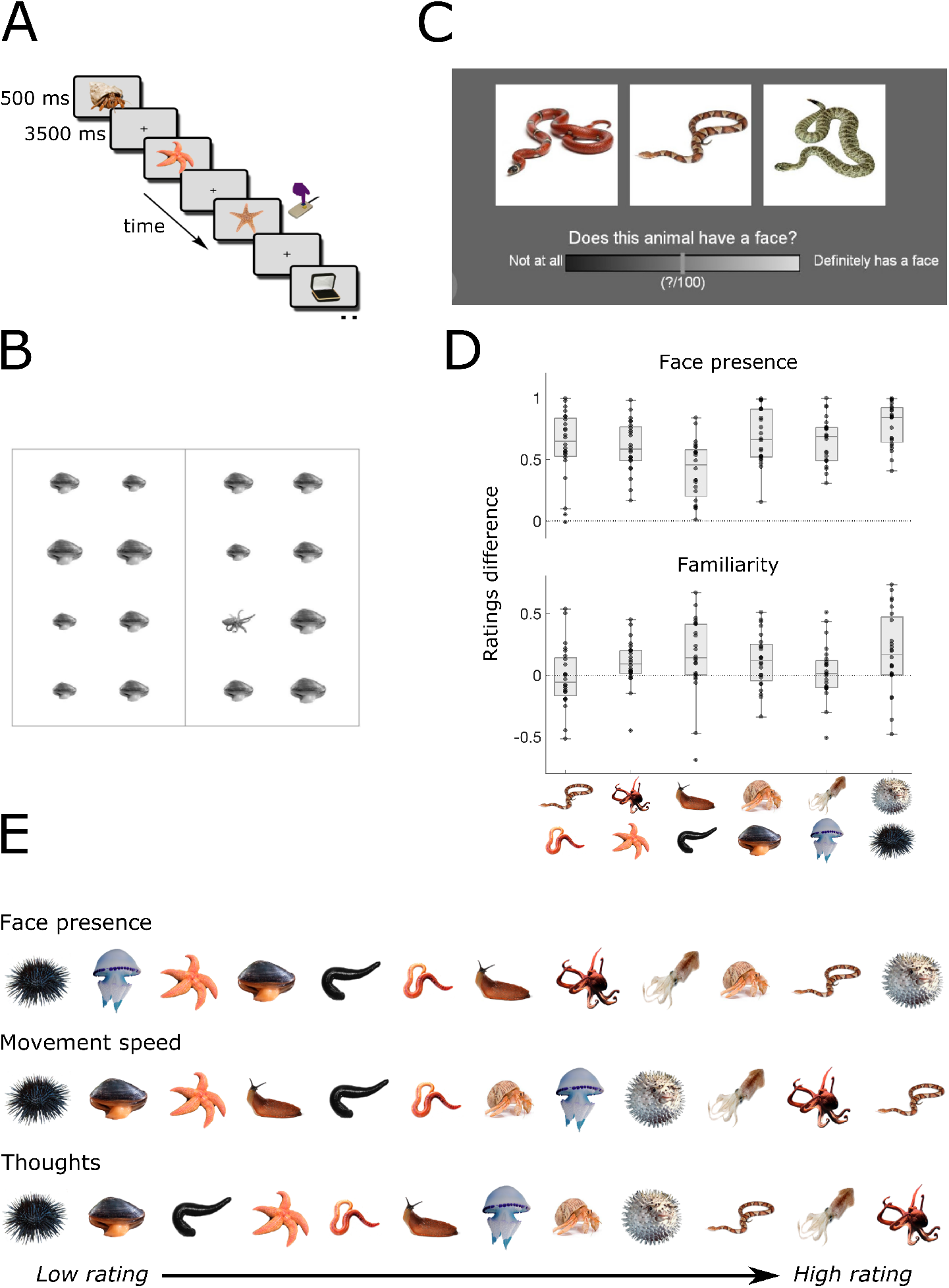
Experimental design and behavioral ratings analysis. ***A***, fMRI experimental design. Stimuli appeared one at a time, and participants were instructed to press a button whenever the same animal or object appeared on 2 consecutive trials. ***B***, Trial example from the behavioral visual search experiment used for quantifying pairwise visual dissimilarity between the stimuli. Participants had to find an oddball image among the distractors as quickly as possible and indicate with a button press whether it was on the left or right of the central line. ***C***, Trial example in a behavioral ratings experiment. Participants had to indicate their response by clicking the mouse at any point of a continuous ratings bar. ***D***, The difference between the face presence (upper panel) and familiarity (lower panel) ratings for each of the 6 animal pairs. Above zero values indicate that, for the 2 animals sharing a similar shape, the animal with a face received higher rating than an animal without a distinct face. ***E***, Animate stimuli sorted based on group-averaged behavioral ratings, from the lowest to the highest.

### Behavioral visual search experiment

The visual search experiment was analogous to the one described in Proklova et al. (2016) and based on the approach by Mohan and Arun (2012). On each trial, participants saw an array of 16 stimuli, 15 of which were identical distractors and one was an oddball target (see Fig. 2B for a trial example). We measured participants’ reaction times (RTs) to find the target and used the RT as a proxy of visual similarity between target and distractor. Faster RTs suggest that target “popped out” more because it was more visually distinct from distractors, and slower RTs indicate that the search was harder because the target was quite similar visually to the distractors. Each of the stimuli (Fig. 1) appeared both as a target and a distractor during the experiment, giving us the visual similarity estimate for each pair of stimuli. This excluded comparisons between the 3 versions of the same animal or object, since the responses for 3 versions were averaged in the fMRI experiment and were not analyzed separately in the representational similarity analysis. We refer the reader to Proklova et al. (2016) for further details of the visual search experiment.

### fMRI acquisition

fMRI data were acquired using a Siemens MAGNETOM Prisma Fit whole-body 3 Tesla MRI scanner with a 32-channel head coil at the Centre for Functional and Metabolic Mapping, Western University, London, Ontario. The stimuli were back projected onto a translucent screen placed inside the scanner bore (60 Hz refresh rate; 1024 × 768 spatial resolution). Participants viewed the stimuli via a mirror mounted to the head coil. Functional images were collected with an echo planar imaging sequence (echo time, 30 ms; repetition time, 2000 ms; field of view, 196 × 196 mm^2^; matrix, 64 × 64; flip angle, 90°; slice thickness, 3 mm; gap, 0 mm; number of slices, 36; axial slice orientation). A high-resolution 3D structural T1-weighted scan was collected at the beginning of the session using a magnetization-prepared rapid gradient echo sequence (MPRAGE) with voxel size of 1 mm isotropic.

### fMRI task

While in the scanner, participants completed 8 runs of the main experiment, and 2 functional localizer runs. In each functional run, each of the 54 stimuli appeared at least once, in randomized order. Some stimuli appeared twice in order to introduce one-back repetition trials (10 per block.) Participants’ task was to press the button any time the image of the same animal or object (e.g., 2 snakes, but not necessarily the exact same photo of a snake) appeared on 2 consecutive trials. The button-press trials were not further analyzed. Stimuli were on screen for 500 ms, followed by 3500-ms inter-stimulus interval (Fig. 2A). Each run lasted 256 s. In the functional localizer runs, participants saw images from 4 categories (faces, bodies, animals, objects) in a blocked design, pressing the button when the same image appeared twice in a row. The localizer procedure is described in detail in Proklova et al. (2016).

### fMRI preprocessing

The neuroimaging data were analyzed using Matlab and SPM12. Preprocessing involved realigning the functional volumes, coregistering them to the structural image, re-sampling to a 2 × 2 × 2 mm grid, and spatially normalizing to the Montreal Neurological Institute 305 template included in SPM12. For the univariate analysis of the localizer data, the functional images were smoothed with a 6-mm FWHM kernel. Images were not smoothed for the multivariate analyses of the main experiment data. For the main experiment, the BOLD signal of each voxel in each participant was modeled using 24 regressors in a general linear model, with 18 regressors for each of the objects (e.g., one regressor for all snakes) and six regressors for the movement parameters obtained from the realignment procedure.

### ROI definition

The four regions of interest (ROIs) are shown in Figure 3A and were defined as follows. Early visual cortex (EVC) ROI was defined anatomically by selecting Brodmann area 17 (BA17) using WFU PickAtlas toolbox for SPM12 (Maldjian et al., 2003), and spanned 5,856 mm3 in size. To define fusiform face area (FFA), we used the group-level Faces > Houses contrast in the functional localizer (p < 0.05, FWE corrected), which revealed two clusters: one in the right hemisphere (792 mm3, peak MNI coordinates: x = 46, y = −50, z = −20), and one in the left hemisphere (208 mm3, peak MNI coordinates: x = −42, y = −40, z = −20). Ventral temporal cortex (VTC) was defined according to previous studies (Haxby et al., 2011, Thorat et al., 2019). It included the inferior temporal, fusiform, and lingual/parahippocampal gyri, and extended from −71 to −21 on the y-axis of the MNI coordinates, with total volume of 73,776 mm3. Finally, we included a region defined in an earlier study (Proklova et al., 2016) in which the animate/inanimate distinction was observed after controlling for visual features (this study involved different stimuli and participants.) We refer to it as “Animacy ROI” for simplicity. It consisted of two clusters, with peaks in right fusiform gyrus (x = 42, y = −60, z = −18; 2,672 mm3) and left fusiform gyrus (x = −44, y = −52, z = −16; 1,968 mm3). The specific analyses that were used to define this region are described in Proklova et al. (2016). The Animacy ROI and the FFA only shared 10 voxels (80 mm3) in common, and both were almost fully encompassed by the VTC: 97% of FFA (122/125 voxels) and 83% of the Animacy ROI (482/580 voxels) intersected with the larger VTC ROI.

**Figure 3.**
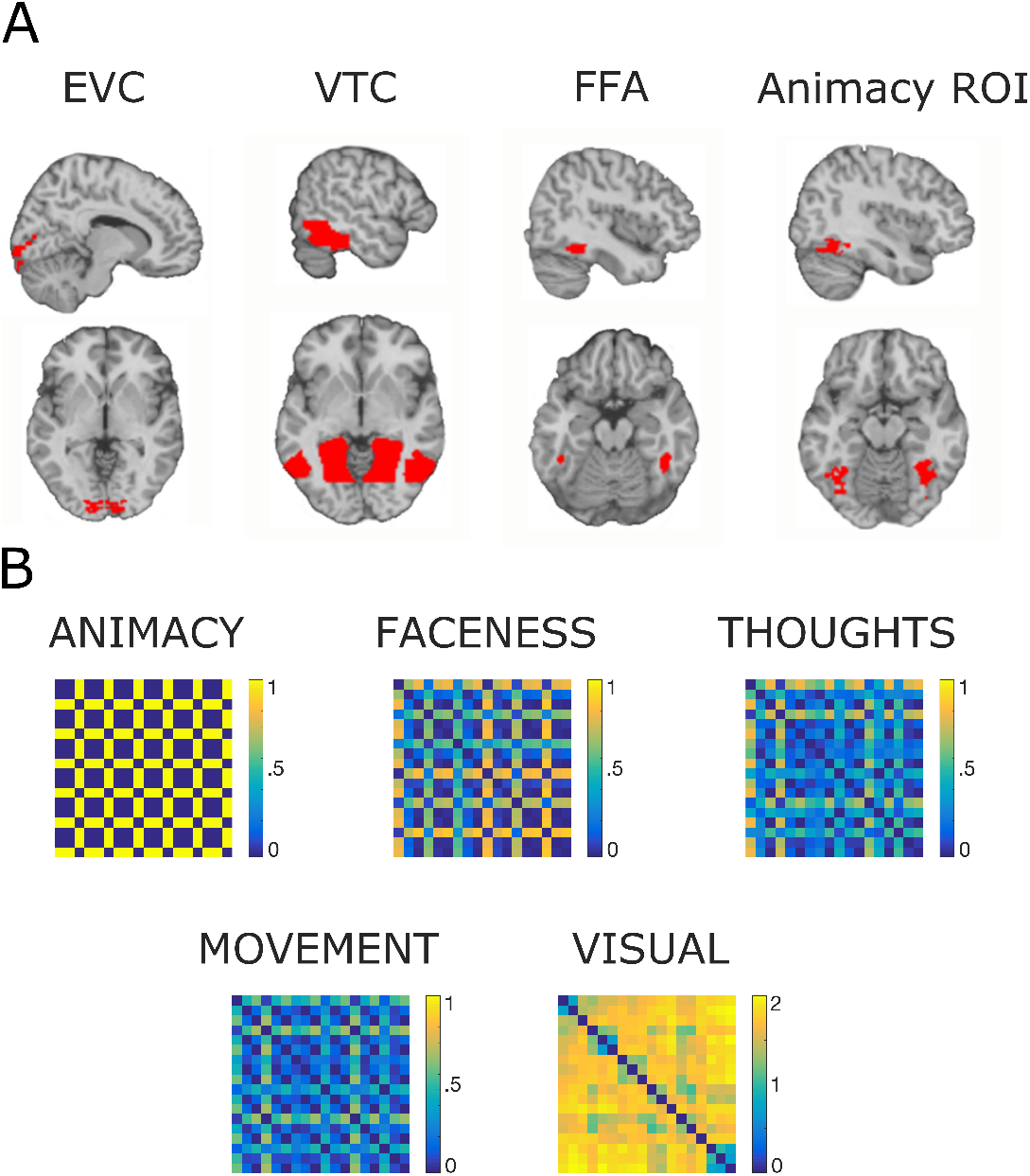
Regions of interest and model RDMs. ***A***, Regions of Interest included early visual cortex, ventral temporal cortex, fusiform face area, and the region in which the animate/inanimate distinction was observed after controlling for visual features in an earlier study (Animacy ROI). ***B***, Model representational dissimilarity matrices (RDMs) used in the representational similarity analysis.

### Representational Similarity Analysis (RSA) Searchlight

The RSA searchlight procedure was analogous to the one used in previous studies (Proklova et al, 2016; Thorat et al., 2020). All analyses were done using CoSMoMVPA toolbox for Matlab (Oosterhof et al., 2016). For each voxel in the brain, we took a 100-voxel spherical neighborhood around it and measured voxel-wise response patterns for all 18 conditions in this region. We then calculated pairwise Pearson correlations between these patterns for all pairs of stimuli. These correlations were then inversed (1-Pearson) and used to create a 18 × 18 neural representational dissimilarity matrix (RDM), in which every entry corresponded to the dissimilarity between a pair of stimuli. For each 100-voxel neighborhood, we then ran a general linear model (GLM) style regression, in which the neural RDM was modeled as a linear combination of two model RDMs: Category (Animacy) and Visual (Fig. 4A), resulting in two beta-weights describing the individual contribution of each model to the neural dissimilarity. Finally, the resulting beta-maps for all participants were entered into a second-level analysis in SPM 12, producing a whole-brain maps reflecting the contributions of Animacy and Visual information to the VTC response patterns (Fig. 4B). Additional details about the Searchlight procedure can be found in Proklova et al., 2016. We also ran two additional versions of the RSA using smaller subsets of a full stimulus set. For example, in order to exclude animals with faces from the analysis, we removed the entries of the neural and model RDMs that corresponded to 6 animals with faces, resulting in smaller 12 × 12 RDMs that included only faceless animals and inanimate objects. The same logic was applied when excluding faceless animals from the analysis. Apart from this, the RSA procedure was identical to the one described above.

**Figure 4.**
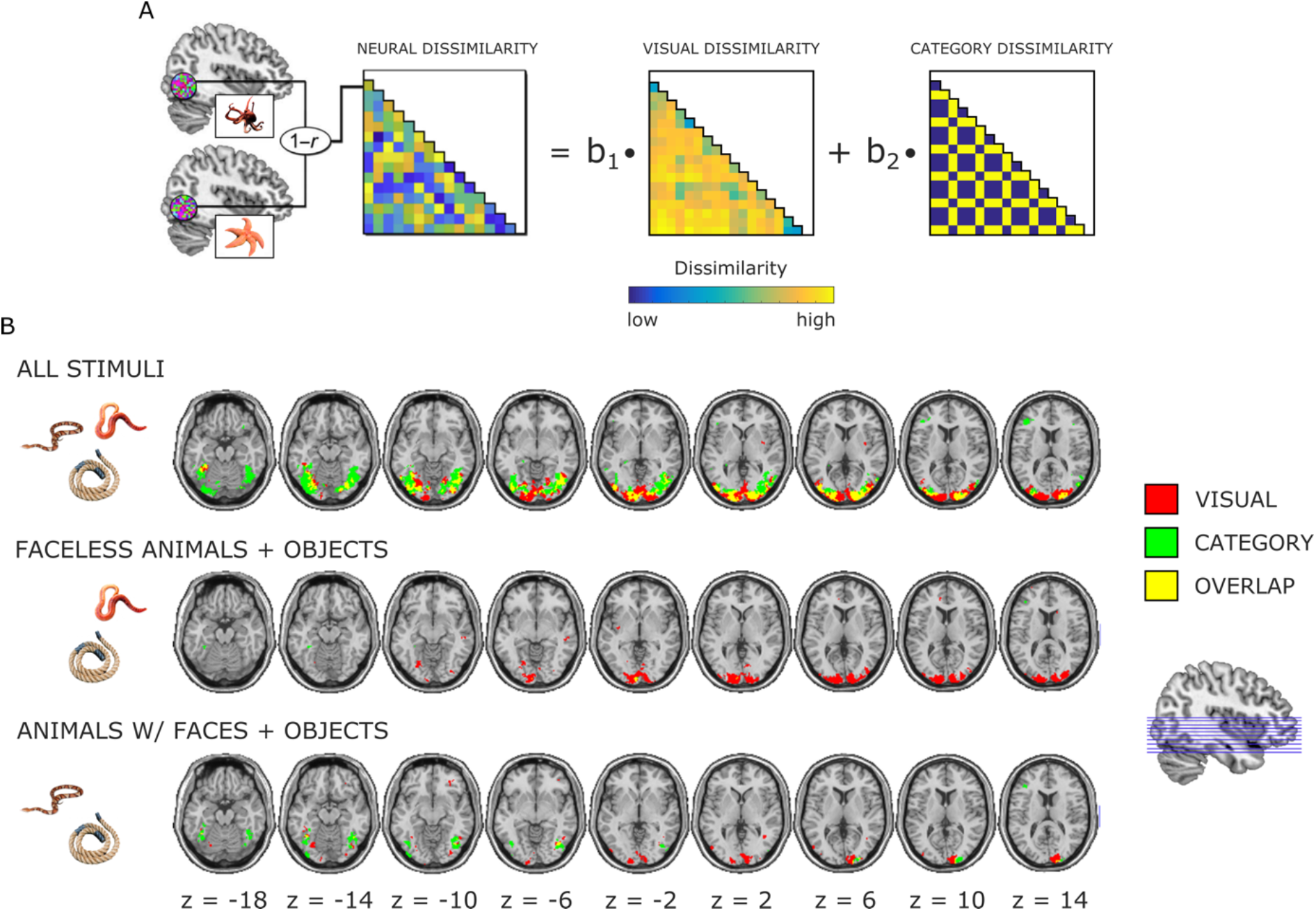
GLM Searchlight. ***A***, Schematic of the analysis. For each voxel, we defined a 100-voxel neighborhood around it and modeled the neural dissimilarity in this region as the linear combination of Visual and Category (Animacy) dissimilarity. ***B***, Searchlight results. Results of whole-brain group-averaged analyses testing the value of each predictor versus zero. The results show independent contributions of Visual (in red) and Category (in green) predictors to neural data. This analysis was run three times: first, including all the stimuli (upper row), next, after removing animals with faces (middle row), and, finally, including only animals with faces and inanimate objects were included (bottom row).

### ROI-based RSA

In the ROI-based RSA, the pairwise neural dissimilarity was measured in the same way as in Searchlight RSA described above, but instead of doing it for each voxel in the brain, it was done in each of the four ROIs (Fig. 3A). After constructing the neural RDMs, we then correlated them with 5 model RDMs that corresponded to Animacy, Face presence, Movement speed, Thoughtfulness, and Visual dissimilarity (Fig. 3B). The neural and model RDMs were normalized before running this analysis. The correlation values were t-tested against zero, and the resulting p-values were corrected for multiple comparisons (Bonferroni correction, 3 analyses × 4 ROIs × 5 correlations = 60 tests, adjusted alpha threshold 0.05/60 = 0.0008). The Animacy RDM was defined by assigning 1 (maximum dissimilarity) to pairs of stimuli belonging to the same category (animate or inanimate), and 0 (minimum dissimilarity) to pairs from different categories. The Faceness, Thoughts and Movement RDMs were defined based on the behavioral ratings from the ratings experiment described above, correlating the ratings for each pair of images. Inanimate objects (not included in the ratings experiment) were assigned a rating of zero. Finally, the Visual RDM was constructed using the reaction times from the behavioral visual search task described earlier. For each pair of stimuli, we used the inverse reaction time (1/RT) for the corresponding target-distractor pair as a corresponding entry to the visual RDM. Longer reaction times (indicating higher visual similarity) were thus reflecting lower visual dissimilarity.

### Multidimensional scaling

To visualize the relationship between the stimuli representations in each ROI, we performed multidimensional scaling (MDS) using the *cmdscale* function in Matlab r2018b, The MathWorks, Natick, MA.

## Results

### Behavioral results

We first wanted to check if our pre-selected faceless animals were indeed perceived as faceless by participants. In our design, each of the six faceless animals (e.g., a worm) was matched with a similarly shaped animal that had a more distinct face (e.g., a snake). For each participant, we took the difference between the two “face presence” ratings for each stimulus pair. The positive difference meant that an animal with a face received a higher “face presence” rating than a corresponding faceless animal. As seen in the upper panel of Figure 2D, this was the case for all six animal pairs. By contrast, there was no significant difference in familiarity between animals with and without faces for each of the six stimulus pairs (Fig. 2D, lower panel).

Next, we averaged the ratings across participants and arranged the animal stimuli on a scale from the lowest to highest “face presence” rating (Fig. 2E), revealing that, on average, all of the six preselected faceless animals were indeed rated lower on the “faceness” scale than the six animals with faces (p < 0.0001). The same analysis was performed with the ratings of movement speed and capacity for thought (see Fig. 2E). On average, animals with faces were rated as faster moving and more capable of thought compared to faceless animals (p = 0.02 and p = 0.002, respectively.) The ratings of head and eye presence were almost identical to face presence ratings and were not further analysed.

Perhaps unsurprisingly, familiarity ratings correlated positively with face presence (r = 0.42), movement speed (r = 0.64), and thoughtfulness (r = 0.54). It can be challenging to disentangle familiarity from these factors because we tend to be more familiar with animals that move and think. As mentioned above, however, there was no significant difference in familiarity between animals from the same shape pair with and without a face.

### RSA Searchlight Results

Our next goal was to replicate the animate/inanimate distinction in VTC representations found in earlier studies (e.g. Proklova et al, 2016) and to explore the possibility that these results could have been driven by animal faces.

We performed the representational similarity searchlight analysis (identical to the one used in Proklova et al, 2016) to reveal independent contributions of Animacy and Visual models to object representations on the whole brain level (Fig. 4, see Methods for the description of the analysis.) Importantly, this analysis was run three times: once with all stimuli (Fig. 4B, upper row), once with only faceless animals and inanimate objects (Fig. 4B, middle row), and once only with animals with faces and inanimate objects (Fig. 4B, bottom row). This allowed us to directly examine the contribution of animal faces to the animate/inanimate distinction in the brain.

The first analysis, which included all the stimuli, revealed bilateral clusters in the VTC in which Animacy model correlated significantly with the neural RDM (Fig. 4B, upper row), with local peaks in left fusiform gyrus (19,096 mm3, peak coordinates: x = −38, y = −62, z = −14) and right fusiform gyrus (22,624 mm3, peak coordinates: x = 40, y = −58, z = −14). However, when animals with faces were excluded from the analysis, we observed only a small cluster in the left fusiform gyrus (168 mm3, peak coordinates: x = −36, y = − 44, z = −16) in which the neural dissimilarity correlated significantly with category (animacy) dissimilarity (Fig. 4B, middle row). As fewer stimuli (and fewer trials) were included in this analysis, there is a possibility that the reduced animacy-related information was observed due to reduced power. To control for this possibility, we then ran the final searchlight analysis in which animals with faces were included in the analysis and faceless animals excluded, which involved the same number of trials as the previous analysis (Fig. 4B, bottom row). The results showed that the animacy information was again robust in the VTC in two clusters with peaks in left inferior temporal cortex (1,992 mm3, peak coordinates: x = −48, y = −64, z = −10) and right fusiform gyrus (3,336 mm3, peak coordinates: x = 32, y = −72, z = −18). This suggests that the reduced animacy-related response observed in absence of animals with faces was not due to reduced power, but specifically to the absence of a face. Together, these results show that including animals with faces in the analysis leads to much more robust animate/inanimate distinction in the VTC. They also raise a possibility that the animacy information reported in previous studies (e.g., Proklova et al., 2016) could largely be an artefact of faces in the animate stimuli.

### ROI-based RSA Results

The searchlight analysis showed that animal faces clearly play an important role in the VTC representations. What is it about face presence that is driving this effect, and is it different in different sub-regions of the VTC? Since face presence is associated with many different factors (from visual features to perceived intelligence and similarity to humans) we wanted to examine this further, including new behavioral models that captured different aspects of the stimuli. We also focused on four regions of interest (ROIs) that were defined prior to and independent from the searchlight analysis, including a large VTC ROI, the fusiform face area (FFA), a region sensitive to animate/inanimate distinction independently of visual features (Animacy ROI, see Methods for the details of how it was defined), and early visual cortex (EVC) as a control region.

For each ROI, we correlated the neural representational dissimilarity matrix (RDM) with five model RDMs characterizing Animacy, Faceness (face presence), Movement Speed, Thoughts, and Visual information. The correlation values were then tested against zero, and p-values were corrected for multiple comparisons (Bonferroni correction). The results are shown in Figure 5. Analogously to the searchlight analysis, the ROI-based RSA was run 3 times using different subsets of stimuli in order to directly examine the effect of animal faces on object representations in those regions.

**Figure 5.**
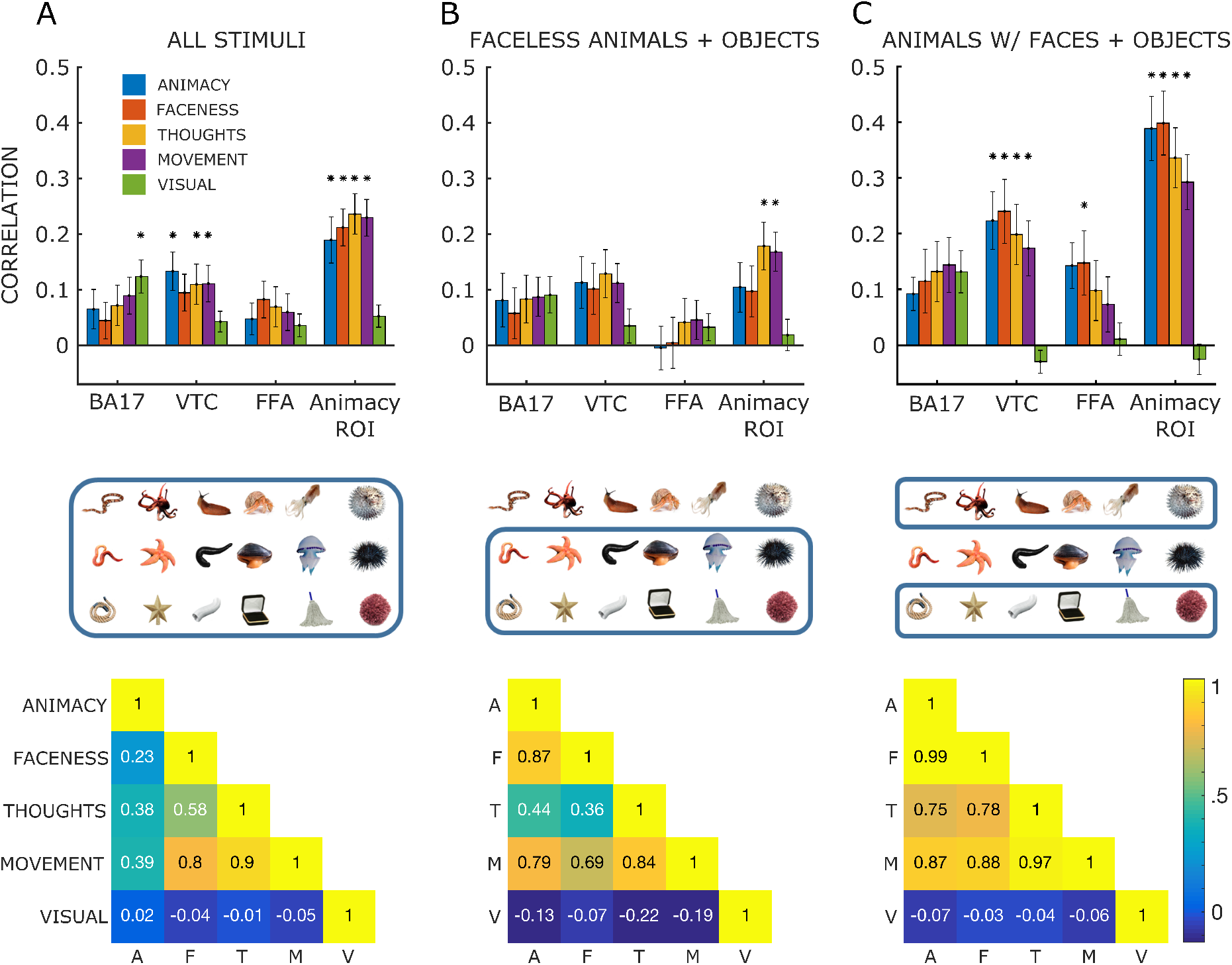
ROI RSA results. Top row: RSA results in four ROIs including all the stimuli ***(A)***, the same analysis repeated after removing animals with faces from the analysis ***(B)***, and after animals with faces were included and faceless animals excluded from the analysis ***(C)***. Asterisks indicate significance after correcting for multiple comparisons. Middle row: stimuli that were included in each type of analysis (in blue frame). Bottom row: pairwise correlations between model representational dissimilarity matrices (RDMs) used in the three analyses.

We first ran this analysis with the full stimulus set (Fig. 5, left column). As expected, the Visual model, but not the high-level ones, correlated significantly with the early visual cortex RDM. In the VTC, the Animacy model showed the highest correlation with the neural RDM, followed by the Movement and Thoughts models for which the correlations were also significant. Interestingly, the correlation with the Faceness RDM did not survive correction for multiple comparisons. In the FFA, although none of the correlations survived multiple comparisons correction, the highest correlation was with the Faceness model. Finally, in the Animacy ROI, all high-level models (apart from Visual) correlated significantly with the neural RDM.

Next, to see whether these results were driven by animals with faces, we re-ran the RSA after excluding all animals with faces from the analysis (Fig. 5, middle column). The results showed overall lower correlation values in all the ROIs, and, crucially, animacy information was not significant after correcting for multiple comparisons in both VTC and Animacy ROI. Interestingly, even in absence of faces, both Movement and Thoughts RDMs still correlated significantly with the neural RDM in Animacy ROI.

Finally, we re-ran the analysis after removing faceless animals and looking only at animals with faces and inanimate objects (Fig. 5, right column). Note that Animacy and Faceness RDMs were almost perfectly correlated in this condition. Strikingly, this led to much higher correlations in VTC and the Animacy ROI with the Animacy RDM and all the other high-level models.

Using different subsets of stimuli in the three analyses meant that the target RDMs and correlations between them also changed. Pairwise correlations between target RDMs are shown in the bottom row of Figure 5. In all three analyses, the Visual RDM did not correlate highly with the remaining high-level models. The correlation between Animacy and Faceness models was 0.23 for the full stimulus set, increased to 0.87 when animals with faces were excluded (likely driven by the fact that in both models the inanimate objects had a rating of zero), and was close to perfect (0.99) in the final analysis when only animals with faces and inanimate objects were included.

At first glance, these results, in line with the Searchlight, seem to suggest that faces heavily influence the representations of animacy in these regions: when animals with faces are excluded, animacy information in VTC is not significant, and when they are present, it is strongly pronounced. If this were the case, however, we would expect to see higher correlations between the Faceness model and the VTC RDM, which was not the case. Instead, this result seems to be driven by something other than faces (but something that correlates with the presence of a face) – in this case, movement speed and capacity to think. Indeed, even when animals with faces were excluded, the Movement and Thoughts models (but not the binary Animacy model) correlated significantly with the neural RDM in the Animacy ROI. This suggests that the animate/inanimate distinction in this region is influenced not so much by the presence of faces, but rather by other properties that correlate with face presence, such as the capacity to move and think.

### Multidimensional scaling results

Finally, we performed multidimensional scaling to visualize the representational structure in each ROI (Fig. 6). In this analysis, the images that are represented similarly in a given ROI end up closer together on a 2D plane. As expected, the representations in the EVC did not show clustering based on Animacy (compared to the other ROIs) and instead seemed to have reflected visual properties: elongated stimuli, such as the tube, slug, and leech, clustering together on the left and roundish objects clustered on the right. In both the VTC and the Animacy ROI, however, the animate/inanimate distinction was pronounced, with inanimate objects clustering together separately from animals (Fig. 6). Intriguingly, this analysis revealed a kind of gradient in those particular ROIs: animals with faces on one side, inanimate objects on the other side, and faceless animals in between. This explains the results of the RSA, showing how including only animals with faces in the analysis makes the distinction between animate and inanimate objects in those regions more pronounced.

**Figure 6.**
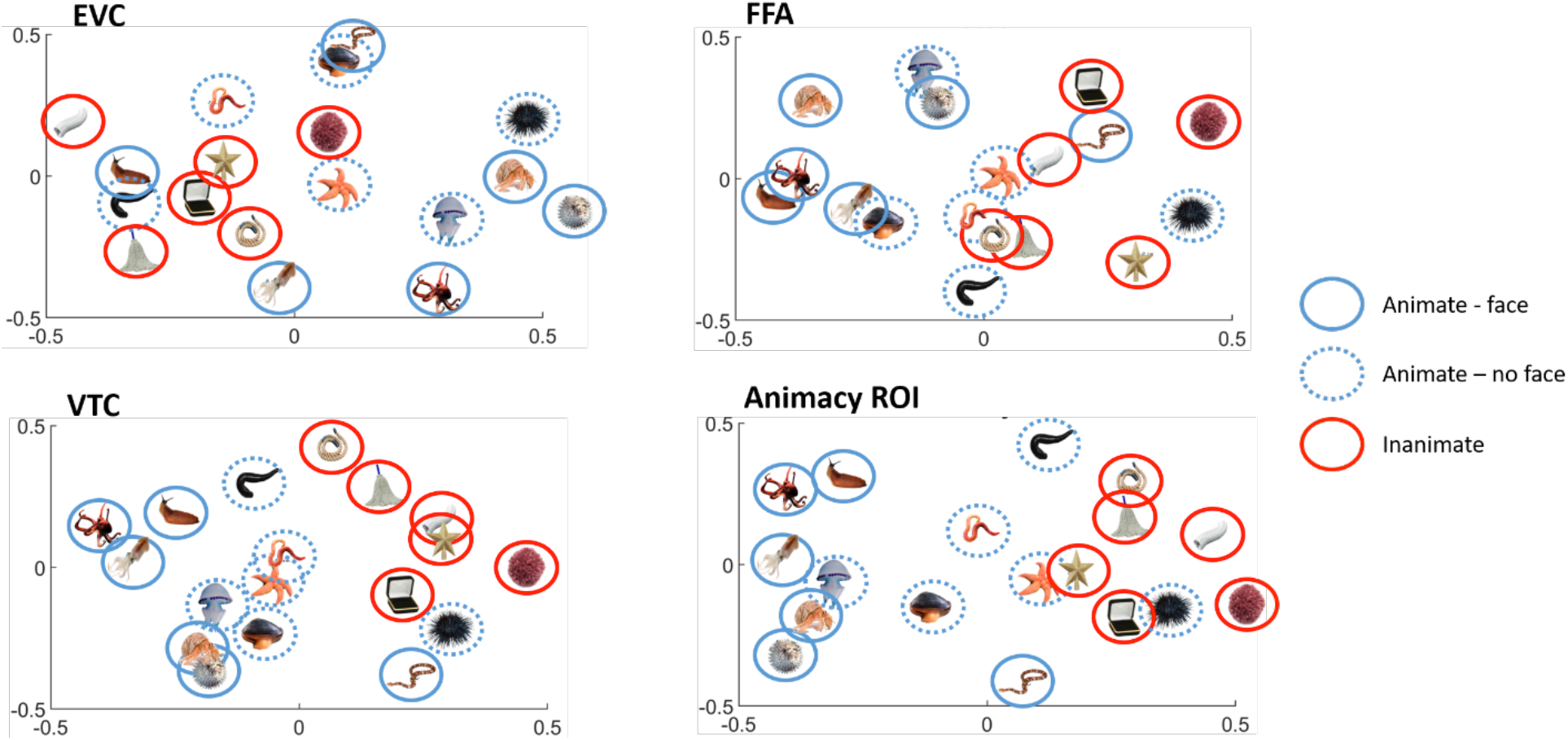
Multidimensional scaling results. Representational structure in the four ROIs revealed by multidimensional scaling, showing how animals with faces (solid blue circles), faceless animals (dashed blue circles), and inanimate objects (red circles) are represented with respect to each other in each ROI. Stimuli that are represented similarly in a given ROI are shown close to each other in 2-dimensional space.

## Discussion

We investigated the contribution of animal faces to the animate-inanimate distinction that has been revealed in ventral temporal cortex (VTC) across many studies (e.g., Kriegeskorte et al., 2008; Grill-Spector and Weiner, 2014; Bracci and Op de Beeck, 2016; Proklova et al., 2016). Unlike previous studies, we systematically controlled for face presence using images of real animals, half of which had a face and half of which did not, as opposed to obscuring the face or using different viewpoints in which a face is turned away (in which case our knowledge of how an animal usually looks could lead to filling-in effects). The initial searchlight analysis revealed that when animals with faces were removed from the analysis, the animate/inanimate distinction almost disappeared. However, further ROI-based representational similarity analysis revealed that Movement and Thoughts models significantly correlated with activity in a sub-region of the VTC – even after faces were removed. Together, these results suggest that the animate/inanimate distinction in the VTC is largely driven not by the presence of faces or animacy *per se*, but rather by perceived agency (a combination of the ability to move and the ability to think) that is correlated with these factors.

### Visual vs. Conceptual Features

How do our findings relate to the visual vs. conceptual features debate about the nature of the animate/inanimate distinction in VTC object representations (Peelen & Downing, 2018; Bracci et al., 2019)? Although we did not address this question directly, our findings do shed some light on the issue. Faces are both a visual and a conceptual feature. They have characteristic visual features, such as eyes and mouth, and even simplistic smiley faces and two dots above a line will elicit a response in the fusiform face area (Caldara et al., 2006; Kim et al., 2016). Seeing faces in inanimate objects, such as clouds, mountains, and tree trunks, is a common experience, underscoring the important biological function of face recognition (Wardle et al., 2020). At the same time, faces convey rich conceptual information, including similarity to humans (Sha et al., 2015; Contini et al., 2019), emotion, and, in case of human faces, information about gender, race, and age (Dobs et al, 2019). Faces are a powerful cue to whether something is animate and, potentially, possessing agency and intelligence. Our findings suggest that VTC processes not just the visual appearance of a face, but also higher-level information for which faces are a proxy: specifically, animal’s capacity for agency (movement and thought). Moreover, we found that animal faces are not necessary for eliciting animate/inanimate distinction in the VTC, in line with earlier studies (Chao et al, 1999; Martin & Weisberg, 2003; Long et al., 2018).

It could still be the case that the animacy-related activity observed when animals with faces were excluded from the analysis was driven by some remaining visual features, such as curvature, symmetry, and visual texture, that differentiated the animals without faces from the inanimate objects. That explanation is unlikely to be the whole story, however, since we selected our images in a way that minimized shape and texture differences, which was confirmed by low correlation between Visual and Animacy models. Moreover, in a previous study (Proklova et al., 2016), we showed that visual features such as overall shape and texture are not driving the animate/inanimate distinction in the VTC. Of course, we still have to rely on visual information in order to recognize faceless animals as animals, but the fact that we observed high-level information about movement speed and capacity for thought in the Animacy ROI strongly suggests that our conceptual knowledge about an animal also comes into play in this region. Many of us have learned through the media and real-life experiences that starfish, sea urchins, and other creatures that at first could appear as inanimate are in fact animals. In other words, semantic associations between the image and previous knowledge is likely driving the observed activation in the animacy-sensitive regions when images of animals without faces are presented.

### A Gradient vs. a Dichotomy

More and more studies show that a simple animate/inanimate dichotomy is not the best way to explain the VTC representational geometry (Bracci et al., 2018; Contini et al., 2020; Connolly et al., 2012; Sha et al., 2016; Carlson et al., 2013). Our findings also suggest that animacy in the VTC is not all-or-none, but graded: from animals that are perceived as more mobile and intelligent to animals that are perceived as less capable of movement and thought (and thus more similar to inanimate objects). Other recent findings have also found that agency is an important organizing principle for the VTC object representations (Thorat et al., 2019; Haxby et al., 2020).

It has also been proposed that this continuum is driven by similarity to humans (Connolly et al., 2012; Sha et al., 2016; Ritchie et al., 2020), which could explain why inanimate objects such as robots and toys are represented similarly to animals in VTC (Bracci et al., 2019; Contini et al., 2020). Our study did not address this directly, since by design all the animate stimuli were quite dissimilar to humans. Our findings do point, however, to the importance of perceived agency (a combination of ability to move and intelligence) for object representations in VTC. Animals (or animal-like objects) that are perceived as possessing agency are indeed more similar to humans, compared to animals that do not move and have simpler nervous systems. That said, our results suggest that an animal does not have to share visual features with humans or to be evolutionary “closer” to them in order to be represented distinctly from inanimate objects in VTC. Capacity to move and intelligence are very behaviorally relevant features when it comes to perception and recognition of animals, regardless of how similar an animal is to a human. After all, a snake shares few physical features with humans, but it is important to recognize it as animate in order to avoid danger – and movement (as well as the face presence) is an important cue.

### Static vs moving stimuli

Like most studies that have explored the distinction between activity related to animate and inanimate objects in the VTC, we presented our participants with static images. Had we used video displays of animals vs. non-animals, then the presence of self-movement or agency certainly would have been a powerful cue to animacy (Martin and Weisberg, 2003; Haxby et al., 2020). In other words, self-movement could be as potent a cue for animacy as faces. The few studies that have used stimuli that move like animals have found a characteristic animate/inanimate distinction in the VTC (Martin and Weisberg, 2003). Moreover, even though we used only static images in our study, the presence of faces in some of the images could easily invoke the concept of movement (and other features associated with animacy). Thus, as we have already discussed, this nexus of associated animacy features could explain why images of animals with faces are represented as more “animate” in the VTC compared to images of faceless animals, thus eliciting a stronger animate/inanimate distinction.

### Implications for Future Investigations

Our study does not speak to how and where the associations between faces and other aspects of animacy are encoded. There is a possibility that high-level aspects of animacy (e.g., agency) are first processed outside of VTC, and this information is then conveyed back to the VTC via re-entrant pathways. Electrophysiological techniques such as M/EEG could shed light on the time course of this process (Cichy & Oliva, 2020). Our findings also suggest that any attempt to disentangle the factors contributing to the animate-inanimate distinction (or gradient) in VTC should pay close attention to the animals that are used as stimuli. It is critical to include a wide range of animate objects, not just more typical, human-looking mammals. Moreover, this applies, not just to animals, but to any object category that is being investigated. The use of large, diverse, and naturalistic stimuli sets (Hebart et al., 2019, Nastase et al., 2020) is one way forward.

## Author contributions

D.P. and M.A.G. designed research; D.P. performed research; D.P. analyzed data; D.P. drafted the manuscript; D.P. and M.A.G. edited the manuscript.

## Acknowledgements

This research was supported by grants from the Natural Sciences and Engineering Research Council of Canada (RGPIN-2017-04088) and the Canadian Institute for Advanced Research to M.A.G. Authors thank Sushrut Thorat and Giacomo Ariani for their helpful feedback throughout the project.

